# Glucose intolerance induces anxiety-like behaviors independent of obesity and insulin resistance in a novel model of nutritional metabolic stress

**DOI:** 10.1101/2023.08.17.553760

**Authors:** Mohammed Al-Onaizi, Kawthar Braysh, Selma Alkefeef, Dana Altarrah, Shorouk Dannoon, Dalal Alasousi, Hawraa Adel, Mariam Al-Ajmi, Anwar Kandari, Rawan Najem, Rasheeba Nizam, Michayla R Williams, Sumi John, Thangavel Alphonse Thanaraj, Rasheed Ahmad, Heba Al-Hussaini, Fahd Al-Mulla, Fawaz Alzaid

## Abstract

Type 2 diabetes (T2D) and its comorbidities are a major public health concern. In addition to peripheral tissues, T2D also impacts the central nervous system leading to neurocognitive impairments, including memory deficits, anxiety, and depression. The metabolic determinants of these neurocognitive impairments remain unidentified. Here, we used a novel proprietary high-fat diet, in which glucose intolerance precedes weight gain, to decipher the metabolic determinants of neurocognitive affects. We show that this model exhibits anxiety-like behaviors, without eliciting depression nor recognition memory deficits. Long-term feeding leads to weight gain, brain glucose hypometabolism and impaired recognition memory alongside the early onset anxiety-like behavior. Using an established genetic model of T2D (db/db) and of diet-induced obesity we show that additional insulin resistance and obesity are associated with depressive-like behaviors and recognition memory deficits. Our findings indicate that glucose intolerance alone can elicit anxiety-like behaviors. Through this study we also provide a novel nutritional model to characterize the discrete effects of glucose intolerance on cognition, behavior, and the physiology of metabolic disease.

## Introduction

Depression and anxiety are recognized comorbidities of type 2 diabetes (T2D) (Deschenes et al., 2023; Hryhorczuk, Sharma, & Fulton, 2013). The risk of T2D begins with lifestyle factors (e.g., a poor diet high in saturated fat and refined sugars), followed by a loss of glucose tolerance, and insulin resistance. At the stage of insulin resistance, pancreatic beta cells first compensate by increasing insulin secretion, they then decompensate and no longer produce insulin, T2D is declared at this stage. Interestingly, cognitive deficits and psychiatric manifestations can arise even at the early stages of glucose intolerance and insulin resistance (Willmann et al., 2020). Given the evolutive nature of T2D, the interactions between specific metabolic states and different aspects of neurocognitive function merit investigation (Wang, Yan, Du, Tao, & Liu, 2021).

We recently demonstrated that a genetic model of T2D (db/db mice) has severely impaired cognition and anxiety-like behavior (Al-Onaizi et al., 2022). Other studies have shown that diet-induced obesity (DIO) and insulin resistance result in anxiety-like and depressive behaviors in rodents (Leffa et al., 2015; Ogrodnik et al., 2019a). Interestingly, the central effects of high-fat feeding have been reported to be time-dependent, where short-term feeding reduces anxiety, but long-term feeding promotes it (Sweeney, O’Hara, Xu, & Yang, 2017). These time-dependent effects indicate that overnutrition alone is not sufficient to induce undesirable neurocognitive outcomes. Such outcomes may be associated with specific consequences of DIO, highlighting the need to further elucidate which components of dysmetabolism impact psychiatric behaviors.

Widely used genetic and nutritional models of metabolic disease (e.g., db/db, ob/ob, DIO) recapitulate multiple components of metabolic syndrome (Burke et al., 2017; Strissel et al., 2007). Namely, obesity, glucose intolerance, insulin resistance in both genetic and nutritional models, in addition to insulinopenia in the db/db model. These multiple components represented in a single model complicate deciphering which will most impact behavior and cognition (Diaz de Leon-Guerrero et al., 2022). What remains missing are the discrete contributions of glucose intolerance and suboptimal nutrition. Defining the components of dysmetabolism that are major determinants of the psychiatric comorbidities of T2D can help guide therapeutic targeting or define which patient populations are most at risk.

Here, we developed an in-house nutritional model that impacts glucose tolerance, independent of insulin resistance, and of short-term weight gain. Benchmarking against established models, we investigated effects of our in-house nutritional model on cognitive function, anxiety, and depression in mice. We show that glucose intolerance alone is associated with anxiety-like behaviors, but not depressive-like behaviors nor memory deficits. Long-term administration results in weight gain alongside glucose intolerance, impaired recognition memory, and sustained anxiety-like behavior. These specific phenotypes were associated with dysregulated glucose metabolism in the brain. Thus, our nutritional model reveals that glucose intolerance *per se* is sufficient to induce anxiety, whereas concurrent weight gain impacts other cognitive domains. These findings indicate that the plethora of neurocognitive outcomes associated with T2D may be driven by specific features of dysmetabolism.

## Results

### A proprietary high-fat diet rich in saturated fatty acids, mono-unsaturated fatty acids and poor in poly-unsaturated fatty acids influences whole body metabolism

Dietary lipid content and composition, including structural factors like carbon number and saturation, influence the outcomes of high-fat feeding (DeLany, Windhauser, Champagne, & Bray, 2000). One of these outcomes, both important and sensitive to lipid composition, is anxiety-like behavior (Nakajima, Fukasawa, Gotoh, Murakami-Murofushi, & Kunugi, 2020). We developed a proprietary high-fat diet (P-HFD), higher in fat content than a normal chow diet (NCD), to induce moderate metabolic stress (see methods). We began by comparing our P-HFD to a commercially available high-fat diet (C-HFD) commonly applied in metabolic disease research to induce obesity, insulin resistance and anxiety-like behavior (Ogrodnik et al., 2019b). In macronutrient proximate analysis, P-HFD was composed of 17.32% calories from protein, 41.13% calories from carbohydrates and 41.56% calories from fat. For comparison, according to manufacturers, NCD (Ssniff, Germany) is composed of 17.08% calories from protein, 72.45% calories from carbohydrates and 10.48% calories from fat and a C-HFD (Research diets, New Brunswick, NJ) is composed of 20% calories from protein, 20% calories from carbohydrates and 60% calories from fat.

We next analyzed fatty acid composition of the P-HFD by gas chromatography, distribution of saturated, monosaturated and polyunsaturated fatty acids (SFA, MUFA, PUFA) was 32.5% SFA, 53.6% MUFA and 13.9% PUFA in P-HFD. Fatty acid composition for the NCD and C-HFD were retrieved from the manufacturer: 16% SFA, 21.9% MUFA and 62.1% PUFA in NCD; and 32.2% SFA, 35.9% MUFA, 31.9% PUFA in C-HFD (Fig. 1B). Thus, the distinguishing feature of both P-HFD and C-HFD is a much higher proportion of SFAs compared to NCD. When comparing the P-HFD to the C-HFD, we find higher proportions of MUFAs and lower proportions of PUFAs in P-HFD.

**Figure 1.**
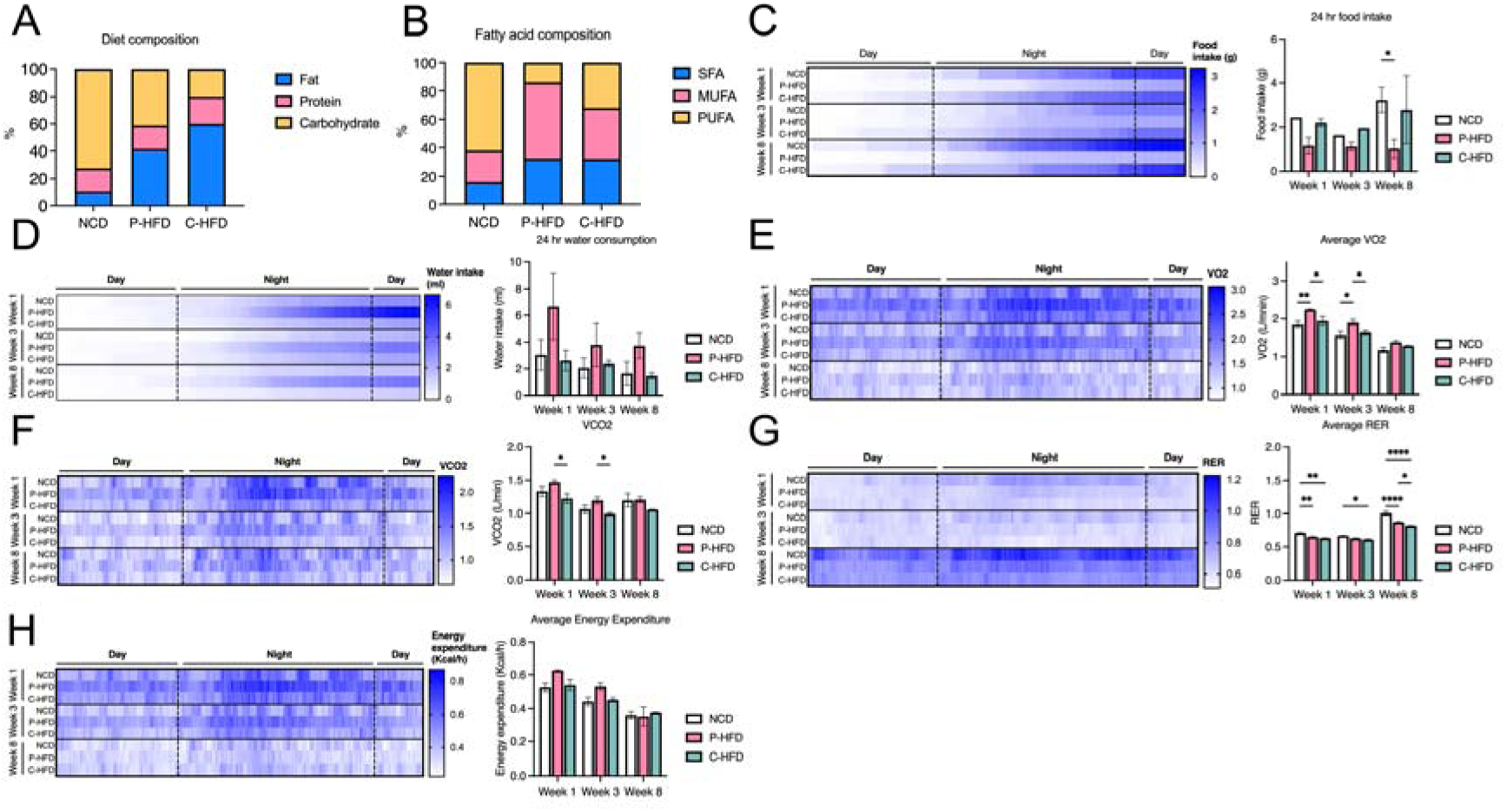
MUFA-rich P-HFD influences whole body metabolism. A. Proximate dietary composition of fat, protein, and carbohydrates in NCD, HFD and C-HFD (expressed as percentage of diet). B. Fatty acid composition of SFA, MUFA, and PUFA in NCD, HFD, and C-HFD (expressed as percentage of diet). C. Heatmap and quantification comparing NCD, P-HFD, and C-HFD groups in 24-hour calorimetric analysis for food intake at week 1, week 3, and week 8 of diet. D. Heatmap and quantification comparing NCD, P-HFD, and C-HFD groups in 24-hour calorimetric analysis for water consumption at week 1, week 3, and week 8 of diet. E. Heatmap and quantification comparing NCD, P-HFD, and C-HFD groups in 24-hour calorimetric analysis for average VO_2_ at week 1, week 3, and week 8 of diet. F. Heatmap and quantification comparing NCD, P-HFD, and C-HFD groups in 24-hour calorimetric analysis for average CO_2_ at week 1, week 3, and week 8 of diet. G. Heatmap and quantification comparing NCD, P-HFD, and C-HFD groups in 24-hour calorimetric analysis for average RER at week 1, week 3, and week 8 of diet. H. Heatmap and quantification comparing NCD, P-HFD, and C-HFD groups in 24-hour calorimetric analysis for average energy expenditure at week 1, week 3, and week 8 of diet. (n=3-4/group, *p<0.05, **p<0.001, ****p<0.0001). SFA, saturated fatty acids, MUFA, monounsaturated fatty acids, PUFA, polyunsaturated fatty acids, RER, respiratory exchange ratio. Data are mean ± SEM.

To characterize the effects of these diets *in vivo* we placed mice on a NCD, P-HFD or C-HFD. We first subjected mice to indirect calorimetry at the beginning of weeks 1, 3, and 8 of feeding to assess food intake, water consumption, gas exchange and energy expenditure. There were no significant differences between NCD, P-HFD, and C-HFD, food intake at 1 and 3 weeks (Two-way ANOVA: no effect of weeks F_(2,12)_=1.02, p=0.3886, an effect of diet, F_(2,12)_=4.91, p=0.0276), and no interaction F_(4,12)_=0.53, p=0.7714. Fig. 1C). Post-hoc analysis revealed that at week 8, P-HFD mice ate significantly less compared to NCD (p=0.0348 Fig. 1C). Calorimetric analysis revealed no significant differences between NCD, P-HFD, and C-HFD for water intake at 1, 3 and 8 weeks (Two-way ANOVA: no effect of weeks F_(2,13)_=0.9965, p=0.3957, no effect of diet F_(2,12)_=2.790, p=0.0981, and no interaction F_(4,13)_=0.2125, p=0.9268, Fig. 1D). In terms of oxygen consumption, average VO_2_ was significantly different between diets (Two-way ANOVA: effect of weeks F_(2,14)_=74.32, p<0.0001, an effect of diet, F_(2,14)_=17.11, p=0.0002, and no interaction F_(4,14)_=0.7814, p=0.557, Fig. 1E). In weeks 1 and 3, post-hoc analysis revealed that P-HFD group had higher VO_2_ compared to NCD (p=0.0030, p=0.0107 respectively), and C-HFD (p=0.0193, p=0.0382 respectively) (Fig. 1E). Average VCO_2_ was also significantly different between diets (Two-way ANOVA: effect of weeks F_(2,14)_=16.83, p=0.0002, effect of diet, F_(2,14)_=10.32, p=0.0018, no interaction F_(4,14)_=0.4960, p=0.7390, Fig. 1F). Post-hoc analysis showed that the P-HFD group had significantly higher VCO_2_, when compared to C-HFD in weeks 1 (p=0.0149) and 3 (p=.0441) (Fig. 1F). The average respiratory exchange ratio (RER) showed significant differences between diets (Two-way ANOVA: effect of weeks F_(2,15)_=416.7, p<0.0001, effect of diet, F_(2,15)_=54.39, p<0.0001, interaction F_(4,15)_=7.216, p=0.0019, Fig. 1G). In week 1, post-hoc analysis revealed that both P-HFD and C-HFD groups had significantly lower average RER, compared to NCD (p=0.0058, p=0.0020, respectively, Fig. 1G). Similarly, in week 3, post-hoc analysis showed significantly lower average RER in P-HFD and C-HFD groups, compared to NCD (p=0.00579, p=0.0188, respectively, Fig. 1G). In week 8, post-hoc analysis showed that the average RER was significantly lower in C-HFD compared to P-HFD (p=0.0150) and NCD (p<0.0001), with P-HFD still lower in average RER compared to NCD (p<0.0001) (Fig. 1G). Energy expenditure data showed no significant differences between the groups across the three timepoints (Two-way ANOVA: effect of weeks F_(2,15)_=20.46, p<0.0001, no effect of diet, F_(2,15)_=2.497, p=0.1159, and no interaction F_(4,15)_=1.030, p=0.4236, Fig. 1H).

Taken together, these data indicate that P-HFD increases metabolism of both fats (VO_2_) and carbohydrates (VCO_2_) during early exposure (weeks 1 to 3), as compared to the NCD and C-HFD. When comparing P-HFD to C-HFD, RER results indicate no difference in predominant energy source between weeks 1 and 3, both of which increase reliance on lipids as compared to NCD.

### P-HFD induces early glucose intolerance but not insulin resistance

Several studies showed that intervention times with high-fat feeding lead to weight gain and a variety of metabolic consequences including glucose intolerance and insulin resistance, ranging from 8 days to 27 weeks (Blancas-Velazquez, Mendoza, Garcia, & la Fleur, 2017; de Moura et al., 2021; Matias et al., 2018; Wu et al., 2018). To determine these kinetics with P-HFD we placed mice on this diet, or a NCD diet. Random glycemia was measured and mice were weighed regularly. Mice were subjected to insulin and glucose tolerance testing (ITT, GTT) at 11-13 weeks, a common timepoint for developing glucose intolerance and insulin resistance on C-HFD, and at 28-32 weeks, an extended time-point for pronounced behavioral effects (Dalmas et al., 2015; Han et al., 2020) (Fig. 2A). Behavioral testing was performed at weeks 13-15, and 28-32 (Fig. 2A).

**Figure 2.**
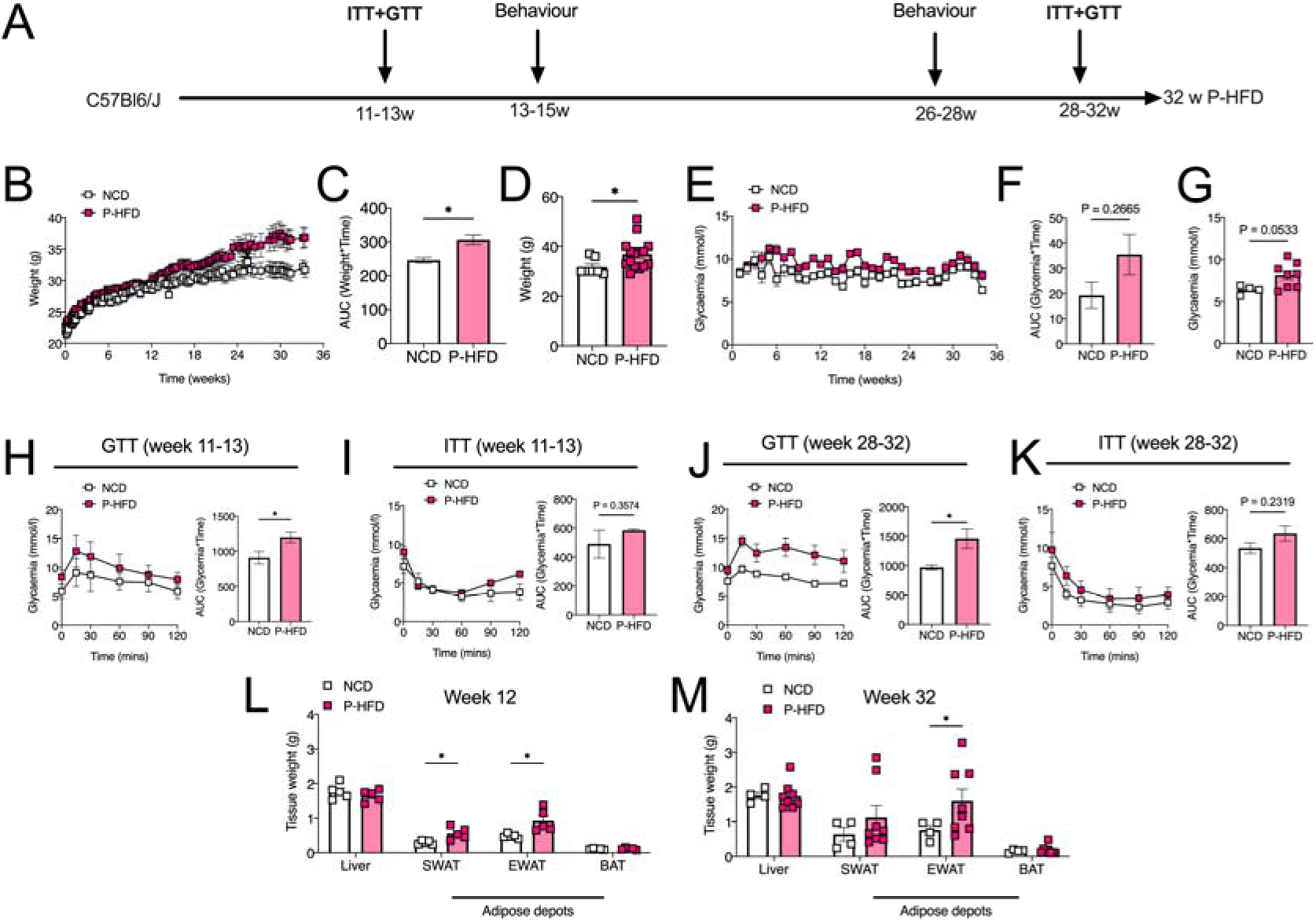
Glucose intolerance precedes weight gain on P-HFD. A. Timeline depicting the duration of P-HFD administration to C56BL6/J mice and metabolic testing (ITT and GTT), and behavioral assessment timepoints. B. Weight analysis across 32 weeks for NCD and P-HFD mice. C. AUC analysis for weight against time for NCD and P-HFD mice. D. End weight analysis at week 32 for NCD and P-HFD mice (NCD, n=9, P-HFD, n=24, *p<0.05). E. Non-fasting glycemia measurements for NCD and P-HFD mice for 32 weeks. F. AUC analysis for non-fasting glycemia levels against time in NCD and P-HFD mice. G. End non-fasting glycemia analysis at week 32 for NCD and P-HFD mice (NCD, n=9, P-HFD, n=24). H. GTT curve for animals at 11-13 weeks of feeding depicting glycemia levels across 120 minutes (left), and AUC analysis (right) for glycemia against time in NCD and P-HFD mice (NCD, n=5, P-HFD, n=5, p<0.05). I. ITT curve for NCD and P-HFD mice at 11-13 weeks of feeding depicting glycemia levels across 120 minutes (left), and AUC analysis (right) for glycemia against time in NCD and P-HFD mice (NCD, n=5, P-HFD, n=5). J. GTT curve for animals at 28-32 weeks of feeding depicting glycemia levels across 120 minutes (left), and AUC analysis (right) for glycemia against time in NCD and P-HFD mice (NCD, n=9, P-HFD, n=9, p<0.05). K. ITT curve for animals at 28-32 weeks of feeding depicting glycemia levels across 120 minutes (left), and AUC analysis (right) for glycemia against time in NCD and P-HFD mice (NCD, n=7, P-HFD, n=15). L. Weight analysis for tissues collected from NCD and P-HFD at 13 weeks (NCD, n=5, P-HFD, n=5, *p<0.05). M. Weight analysis for tissues collected from NCD and P-HFD at 32 weeks (NCD, n=4, P-HFD, n=7, *p<0.05). GTT, glucose tolerance test, ITT, insulin tolerance test, SWAT, subcutaneous white adipose tissue, EWAT, epididymal white adipose tissue, BAT, brown adipose tissue. Data are mean ± SEM.

P-HFD resulted in delayed weight gain, only after week 13, compared to NCD (Two-way ANOVA: effect of weeks F_(95,2566)_=19.93, p<0.0001, effect of diet, F_(1,2566)_=211.5, p<0.0001, no interaction F_(95,2566)_=0.9252, p=0.6822, Fig. 2B). Area under curve (AUC) was increased across the 32 weeks on P-HFD, compared to NCD mice (p<0.05, Fig. 2C), and end-weight at week 32 was approximately 14% higher following P-HFD, compared to NCD (p<0.05, Fig. 2D). Random glycemia readings showed significant overall differences across 32 weeks between P-HFD mice and NCD mice (Two-way ANOVA: effect of weeks F_(32,858)_=3.490, p<0.0001, effect of diet, F_(1,858)_=89.67, p<0.0001, no interaction F_(32,858)_=0.9401, p=0.5639, Fig. 2E). However, the AUC did not show significant differences in random glycemia (p=0.2665, Fig. 2F), and only a strong trend was observed with approximately 21% increase in end-point random glycemia at week 32 of P-HFD compared to NCD (p=0.0533, Fig. 2G).

At weeks 11-13, GTT showed that mice on P-HFD developed mild glucose intolerance compared to NCD, despite little-to-no difference in body weight (Two-way ANOVA: effect of time F_(5,48)_=6.977, p<0.0001, effect of diet, F_(1,48)_=24.49, p<0.0001, no interaction F_(5,48)_=0.4506, p=0.8108, Fig. 2H). AUC was significantly higher in the P-HFD group compared to NCD (p=0.0377, Fig. 2H). Interestingly, P-HFD did not induce significant insulin resistance when ITT was carried out at 13 weeks of diet (Two-way ANOVA: effect of time F_(5,48)_=9.981, p<0.0001, effect of diet, F_(1,48)_=4.716, p=0.0348, no interaction F_(5,48)_=1.191, p=0.3276, Fig. 2I), with AUC showing no significant differences between P-HFD-fed or NCD-fed mice (p=0.3574, Fig. 2I). Long term administration (28-32 weeks) of P-HFD exacerbated glucose intolerance, compared to NCD (Two-way ANOVA: effect of time F_(5,95)_=3.000, p=0.0147, effect of diet, F_(1,95)_=43.65, p<0.0001, no interaction F_(5,95)_=0.6727, p=0.6451, Fig. 2J), with AUC also being higher on P-HFD (p=0.0111, Fig. 2J). At 28-32 weeks of diet, P-HFD mice did not show insulin resistance (Two-way ANOVA: effect of time F_(5,120)_=53.83, p<0.0001, effect of diet, F_(1,120)_=33.71, p<0.0001, and no interaction F_(5,120)_=1.199, p=0.3137, Fig. 2K), with AUC showing no significant differences (p=0.2319, Fig. 2K).

Following early and late time-points, we collected and weighed major metabolic tissues: liver, subcutaneous white adipose tissue (SWAT), epididymal white adipose tissue (EWAT) and brown adipose tissue (BAT). Of these, SWAT and EWAT weight were increased by 13 weeks of P-HFD (both p<0.05), compared to NCD-fed mice (Fig. 2L), and by 32 weeks of diet, EWAT weight remained higher in P-HFD-fed mice compared to NCD (p<0.05, Fig. 2M).

### P-HFD causes mild loss of metabolic homeostasis compared to C-HFD

Following nutritional analysis of P-HFD and *in vivo* testing (Fig. 1 and 2), we carried out side-by-side comparison C-HFD as a validated nutritional model of metabolic disease. We placed mice on P-HFD, C-HFD or NCD for 13 weeks, weighed them regularly and carried out metabolic testing. As previously observed (Fig 2), body weight on P-HFD was not significantly different when compared to NCD-fed mice at week 13 of the regimens. However, C-HFD significantly increased body weight compared to both P-HFD and NCD, the latter consistent with previous reports (Dalmas et al., 2015)(Two-way ANOVA: effect of weeks F_(13,219)_=15.35, p<0.0001, effect of diet, F_(2,219)_=24.87, p<0.0001, no interaction F_(26,219)_=1.505, p=0.0614, Fig. 3A). Post-hoc analysis revealed significant differences between C-HFD and P-HFD starting from week 10 (p=0.0123), until week 13 (p<0.0001) (Fig. 3A). AUC and end-weights confirmed these results, C-HFD mice gained more weight than P-HFD (AUC p=0.0269, end-weight p=0.0099) and NCD mice (AUC p=0.0047, end-weight p=0.0586), with no significant difference between NCD and P-HFD (Fig. 3B and C). Interestingly, fasting glycemia at 6 weeks was higher in P-HFD-fed mice compared to C-HFD (p=0.003) and NCD (p=0.005) (Fig. 3D). However, at 13 weeks, fasting glycemia was similar between mice fed P-HFD or C-HFD (p=0.5573), with both being significantly higher than NCD feeding (P-HFD vs. NCD, p=0.0036, C-HFD vs. NCD, p=0.0005) (Fig. 3E). From this cohort, we subjected C-HFD and P-HFD mice to metabolic testing and found glucose intolerance to be less pronounced in P-HFD-fed mice compared to C-HFD (Two-way ANOVA: effect of time F_(5,60)_=29.96, p=0.003, effect of diet F_(1,60)_=59.00, p<0.0001, interaction F_(5,60)_=3.969, p=0.0036; Fig. 3F), AUC confirmed this finding (p=0.004; Fig. 3G). ITT with a dose adapted to obese animals (0.7U) revealed improved insulin sensitivity upon P-HFD relative to C-HFD (Two-way ANOVA: effect of time F_(5,60)_=30.36, p<0.0001, effect of diet F_(1,60)_=11.54, p=0.0012, no interaction F_(5,60)_=0.6818, p=0.639; Fig. 3H), and this was represented by a similar trend in AUC (p=0.067; Fig. 3I). Upon analysis of tissue weights, we found that white adipose tissue depots (SWAT and EWAT) were significantly lower in weight in P-HFD-fed mice, relative to C-HFD fed mice (p=0.032, p>0.0001, respectively; Fig. 3J). Taken together these data indicate that although glucose homeostasis is disrupted upon P-HFD; metabolic parameters, weight gain and tissue weights remain less affected than in response to C-HFD.

**Figure 3.**
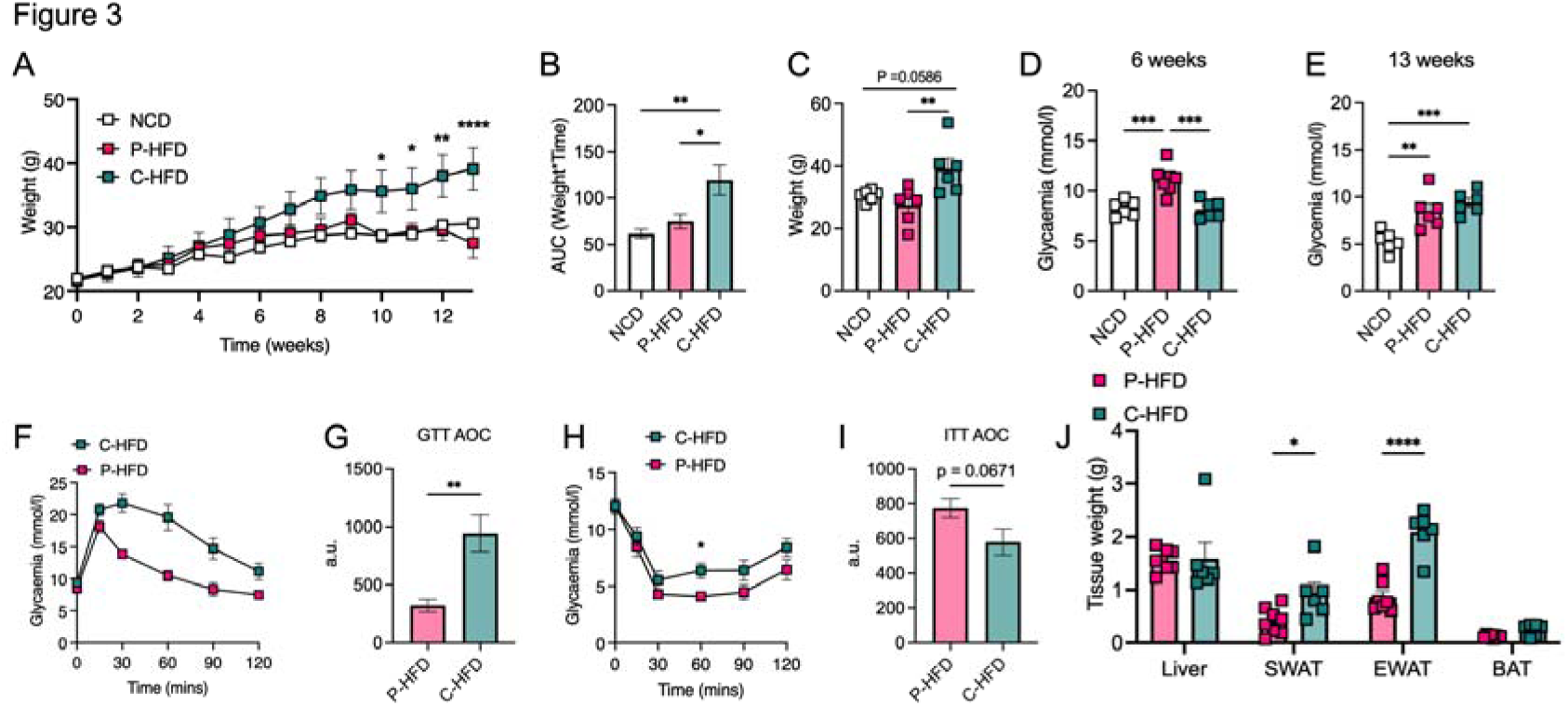
P-HFD causes mild metabolic dysfunction compared to C-HFD. A. Weight analysis across 32 weeks for NCD, P-HFD, and C-HFD mice (NCD, n=6, P-HFD, n=7, C-HFD=6, *p<0.05, **p<0.01, ****p<0.0001). B. AUC analysis for weight against time for NCD, P-HFD, and C-HFD mice (NCD, n=6, P-HFD, n=6, C-HFD=6, *p<0.05, **p<0.01). C. End weight analysis at week 32 for NCD, P-HFD, and C-HFD mice (NCD, n=6, P-HFD, n=6, C-HFD=6, **p<0.01). D. Fasting glycemia levels in NCD, P-HFD, and C-HFD mice at 6 weeks of diet (NCD, n=6, P-HFD, n=7, C-HFD=6, ***p<0.001). E. Fasting glycemia levels in NCD, P-HFD, and C-HFD mice at 13 weeks of diet (NCD, n=6, P-HFD, n=6, C-HFD=6, ***p<0.001). F. GTT curve for P-HFD and C-HFD mice at 13 weeks of diet (P-HFD, n=6, C-HFD=6) G. AUC analysis for GTT curve for P-HFD and C-HFD mice at 13 weeks of feeding. H. ITT curve for P-HFD and C-HFD mice at 13 weeks of feeding depicting glycemia levels across 120 minutes (P-HFD, n=6, C-HFD, n=6, *p<0.05). I. AUC analysis for ITT curve for P-HFD and C-HFD mice at 13 weeks of feeding (P-HFD, n=6, C-HFD=6). J. Weight analysis for tissues collected from NCD and P-HFD at 32 weeks (P-HFD, n=8, C-HFD, n=6, *p<0.05, ****p<0.0001). GTT, glucose tolerance test, ITT, insulin tolerance test, SWAT, subcutaneous white adipose tissue, EWAT, epididymal white adipose tissue, BAT, brown adipose tissue. Data are mean ± SEM.

We investigated specific domains of cognition: learning and memory, anxiety and depressive-like behaviors. To assess memory, we subjected P-HFD, C-HFD at 13 weeks of diet, and db/db mice to the Novel Object Recognition (NOR) task (Fig. 4A). Discrimination index was impaired in C-HFD, and db/db mice compared to controls (One-way ANOVA: effect of groups F_(3,33)_=3.717, p=0.0208, Fig. 4B), with post-hoc analysis revealing significant differences between C-HFD and NCD (p=0.0436), and between db/db and NCD (p=0.0454, Fig. 4B). This is consistent with previous reports (Ogrodnik et al., 2019b; Yermakov et al., 2019). Interestingly, P-HFD-fed mice did not show any impairment in the NOR task compared to NCD, suggesting that P-HFD does not impair recognition and memory, despite loss of glucose tolerance. Moreover, recognition index showed significant differences between NCD and db/db (One-way ANOVA: effect of groups F_(3,32)_=4.443, p=0.0102, post-hoc p=0.0098; Fig. 4C), with post-hoc analysis revealing no differences between NCD and C-HFD (p=0.1022), nor P-HFD (p=0.2746) (Fig. 4C). This suggests impairments in NOR in db/db and C-HFD-fed mice, but not in P-HFD-fed mice.

**Figure 4.**
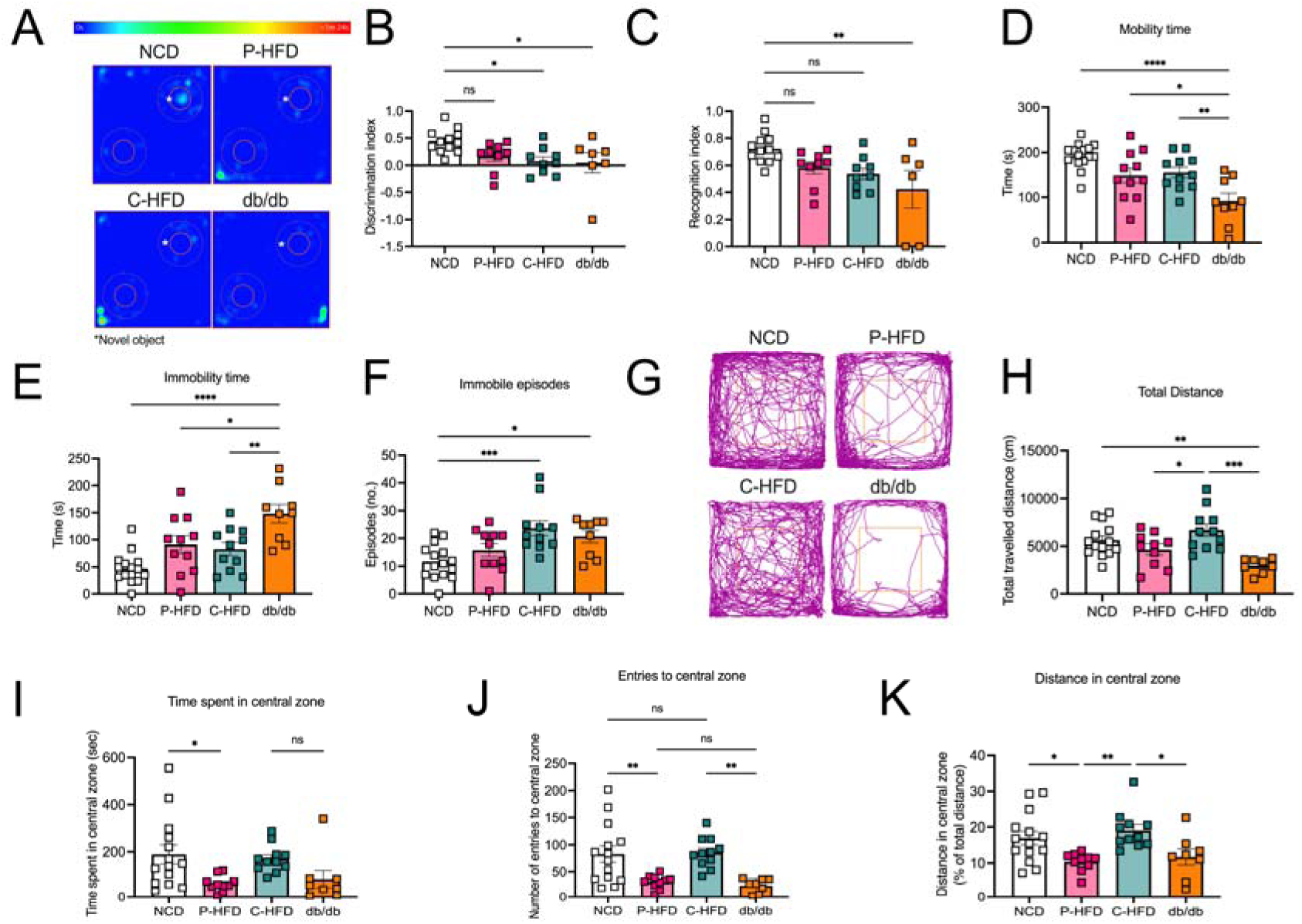
Short-term P-HFD induces anxiety-like behavior, without affecting memory or depression. A. Heatmap representative traces of NCD, P-HFD, C-HFD, and db/db mice on the testing phase of the NOR task depicting time spent investigating the novel object (asterisk). B. Discrimination index of NCD, P-HFD, C-HFD, and db/db mice in the testing phase of the NOR task. C. Recognition index of NCD, P-HFD, C-HFD, and db/db mice in the testing phase of the NOR task (*p<0.05, **p<0.01, NCD, n=12, P-HFD, n=9, C-HFD, n=9, db/db, n=7). C. D. Mobility time of NCD, P-HFD, C-HFD, db/db mice in the FST. E. Immobility time of NCD, P-HFD, C-HFD, and db/db mice in the FST. F. Immobility episodes of NCD, P-HFD, C-HFD, and db/db mice in the FST. (*p<0.05, **p<0.01, ***p<0.001, ****p<0.0001, NCD, n=15, P-HFD, n=11, C-HFD, n=11, db/db, n=9). G. Representative activity traces of NCD, P-HFD, C-HFD and db/db mice on the OFT. H. Total distance travelled in the OFT for NCD, P-HFD, C-HFD, and db/db mice. I. Time spent in the central zone for NCD, P-HFD, C-HFD, and db/db mice in the OFT. J. Number of entries to the central zone for NCD, P-HFD, C-HFD, and db/db mice in the OFT. K. The distance travelled in the central zone for NCD, P-HFD, C-HFD, and db/db mice in the OFT, expressed as percentage of total distance travelled. (*p<0.05, **p<0.01, ***p<0.001, NCD, n=14, P-HFD, n=10, C-HFD, n=10, db/db, n=8). Data are mean ± SEM.

Next, all groups were subjected to the Forced Swim test (FST). Mobility time was significantly different between groups (One-way ANOVA: effect of groups F_(3,42)_=10.64, p<0.0001, Fig. 4D), with post-hoc analysis revealing that db/db mice were less active than all other groups during testing (NCD p<0.0001, P-HFD p=0.0235, C-HFD p=0.0093; Fig. 4D). No overall differences in mobility time were seen between NCD, P-HFD, and C-HFD (Fig. 4D). Similarly, immobility time during the FST showed significant differences across groups (One-way ANOVA: effect of groups F_(3,42)_=10.52, p<0.0001, Fig. 4E), with post-hoc analysis showing that db/db mice were significantly more immobile in the FST compared to NCD (p<0.0001), P-HFD (p=0.0247), and C-HFD mice (p=0.0081) (Fig. 4E). Interestingly, immobility episodes also showed differences across groups (One-way ANOVA shows an effect of groups F_(3,42)_=6.467, p=0.0011, Fig. 4F), with post-hoc analysis revealing that C-HFD-fed and db/db mice had significantly more immobility episodes compared to NCD (p=0.0010, p=0.0283, respectively) (Fig. 4F). These results show that P-HFD-fed mice did not have any impairment in the FST, while db/db mice exhibited overt depressive-like behaviors, and C-HFD displayed moderate depressive-like behaviors.

We then assessed behavior in the Open Field Test (OFT) (Fig. 4G). Data showed differences in the total distance travelled (One-way ANOVA: effect of groups F_(3,39)_=7.742, p=0.0004, Fig. 4H), with post-hoc analysis showing that db/db mice travelled significantly less than C-HFD (p=0.0002), and NCD mice (p=0.0069) (Fig. 4H). Moreover, post-hoc analysis revealed P-HFD-fed mice travelled significantly less than C-HFD-fed mice in the OFT (p=0.0499) (Fig. 4H). To evaluate anxiety-like behavior, we analyzed activity in the central zone of the open field. Only the P-HFD group spent less time in the central zone compared to the NCD group (One-way ANOVA: effect of groups F_(3,39)_=3.936, p=0.0151, Fig. 4I), post-hoc analysis revealed that only P-HFD-fed mice differed from NCD-fed mice, having spent less time in the central zone (p=0.0282). Number of entries to the central zone followed a similar trend, where only P-HFD-fed mice had a lower number of entries compared to NCD-fed mice (One-way ANOVA: effect of groups F_(3,39)_=8.436, p=0.0002; post-hoc p=0.0068; Fig. 4J). Interestingly, the low number of central zone entries by P-HFD-fed mice was at a comparable level to db/db mice that have a much more pronounced metabolic phenotype (Fig. 4J). We then analyzed the distance spent in the central zone as a percentage of the total distance and found significant differences among the groups (One-way ANOVA: effect of groups F_(3,39)_=5.597, p=0.0027, Fig. 4K), with post-hoc analysis revealing that P-HFD-fed mice travelled significantly less in the central zone compared to NCD (p=0.0372), and C-HFD (p=0.0059; Fig. 4K). Moreover, db/db mice travelled a comparable distance to P-HFD-fed mice, and significantly less than C-HFD mice (p=0.0373; Fig. 4K). These findings suggest that P-HFD-fed mice exhibit anxiety-like behavior comparable to db/db mice.

### Long-term P-HFD feeding is associated with memory impairment and anxiety-like behaviors, but not depressive-like behaviors

The above tests demonstrate that short-term glucose intolerance induced by 11-13 weeks of P-HFD triggered anxiety-like behavior, without influencing memory or depression (Fig. 4). These behavioral effects are independent of obesity, insulin resistance and diabetes that is observed in the C-HFD and db/db groups and that influence other behavioral components. Using a separate naïve cohort, we asked whether long-term (32 weeks) P-HFD-feeding that also causes weight gain, but no significant insulin resistance (Fig. 2) would exacerbate specific behavioral outcomes. Contrary to short-term feeding, long-term P-HFD feeding showed robust impairments of discrimination index in the NOR task (p=0.0169, Fig. 5A-B). The P-HFD group also had significant impairments in the recognition index, compared to the NCD group (p=0.0138, Fig. 5C). To determine whether these memory impairments are spatially related we subjected P-HFD-fed mice to the Morris Water Maze (MWM). During the four days of training in the MWM, P-HFD-fed mice were no different to controls in their latency to find the platform (p=0.5900, Fig. 5D). P-HFD-fed mice, however, took less distance to reach the platform during the four days of training, indicating intact learning (Fig. 5E). In the probe trial, no differences were seen between P-HFD and NCD mice (Two-way ANOVA: effect of quadrant F_(1,34)_=104.7, p<0.0001, Fig. 5F), suggesting no impairment in spatial memory in P-HFD mice.

**Figure 5.**
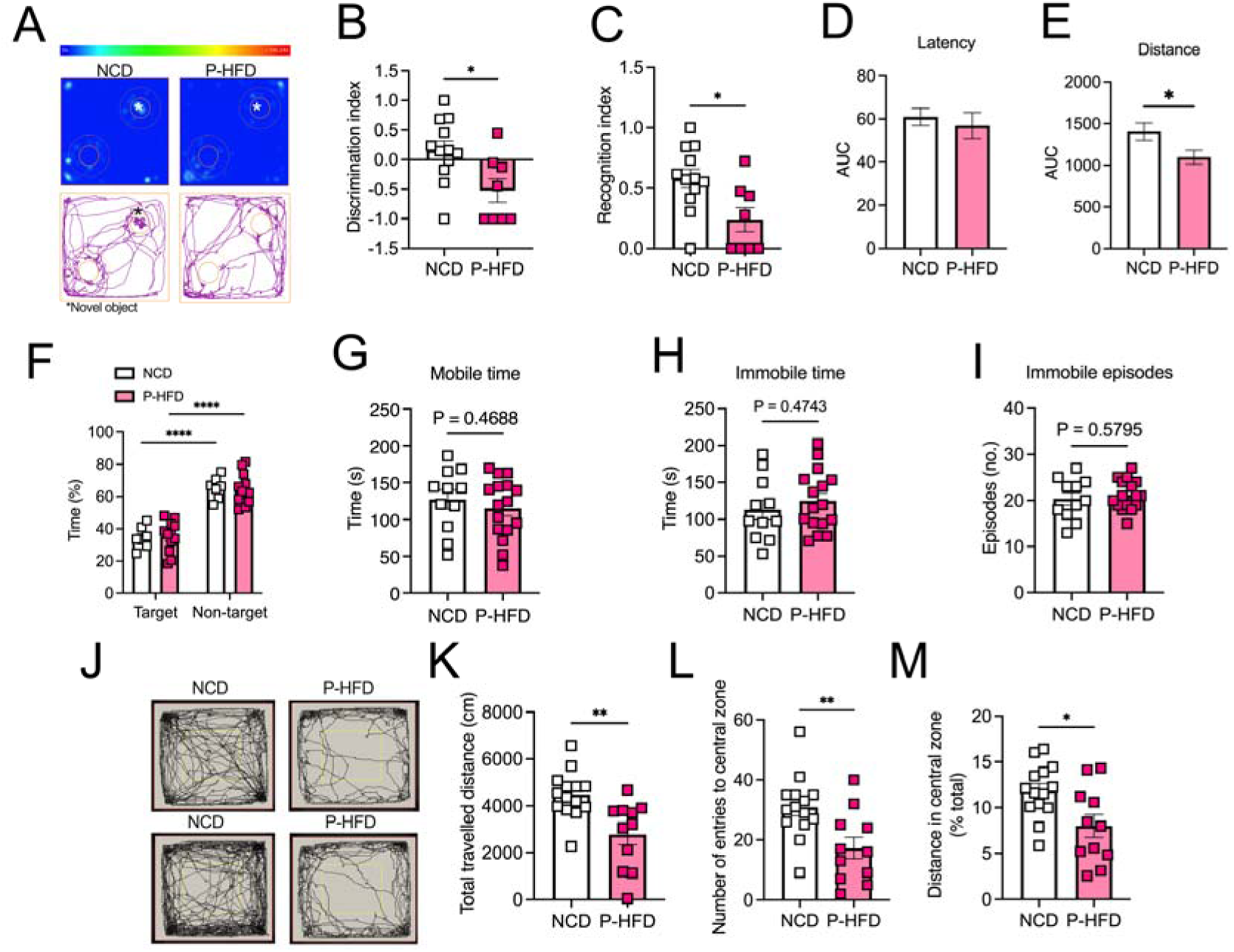
Long-term P-HFD feeding sustains anxiety-like behaviors and impairs memory. A. Heatmap representative traces of NCD and P-HFD on the testing phase of the NOR task depicting time spent investigating the novel object (asterisk). B. Discrimination index P-HFD and NCD mice in the testing phase of the NOR task. C. Recognition index of NCD and P-HFD mice in the testing phase of the NOR task (*p<0.05, NCD, n=12, P-HFD, n=8). D. AUC analysis for latency during the acquisition phase of the MWM task for NCD, and P-HFD mice. E. AUC analysis for distance travelled during the acquisition phase of the MWM task for NCD, and P-HFD mice. F. Percentage time spent in the target vs. the other quadrants in the MWM probe trial for NCD and P-HFD mice (*p<0.05, NCD, n=7, P-HFD, n=12). G. Mobility time of NCD and P-HFD mice in the FST. H. Immobility time of NCD and P-HFD mice in the FST. I. Immobility episodes of NCD and P-HFD mice in the FST (NCD, n=11, P-HFD, n=16). J. N. Representative activity traces of NCD and mice on the OFT. K. Total distance travelled in the OFT for NCD and P-HFD mice. L. Number of entries to the central zone for NCD and P-HFD mice in the OFT. M. The distance travelled in the central zone for NCD and P-HFD mice in the OFT, expressed as percentage of total distance travelled. N. Time spent in the central zone for NC and P-HFD mice in the OFT (*p<0.05, **p<0.01, NCD, n=14, P-HFD, n=11). Data are mean ± SEM.

We then assessed whether long-term P-HFD feeding affected depressive-like behavior and found no significant differences between P-HFD and NCD mice in mobility (p=0.4688, Fig. 5G) nor immobility time in the FST (p=0.4743, Fig. 5H). Similarly, there were no differences between P-HFD and NCD in the number of immobility episodes in the FST (p=0.5795, Fig. 5I), suggesting no depressive-like behavior.

Next, we tested anxiety-like behaviors using the OFT (Fig 5J). Our data showed that long-term P-HFD-fed mice travelled significantly less in the OFT compared to NCD (p=0.0032, Fig. 5K). Analysis of navigation in the central zone of the OFT revealed that P-HFD had significantly less entries to the central zone (p=0.0062, Fig. 5L), and distance travelled in the central zone as a percentage of the total distance travelled during the task (p=0.0119, Fig. 5M). These data suggest that long-term feeding with the P-HFD significantly impairs memory in the NOR task and sustains the anxiety-like behavior seen following short-term feeding, while having no impact on depressive-like behaviors.

### P-HFD feeding leads to glucose hypometabolism in the brain and altered transcriptomic features

Our data demonstrated that glucose intolerance and weight gain, induced by P-HFD feeding, selectively impaired specific cognitive domains in a temporal manner and induced anxiety-like behaviors. To determine consequences on brain health and metabolism, we used glucose analog ^18^F-fluorodeoxyglucose (FDG-PET) imaging in 32-week P-HFD fed mice. We administered ^18^FDG retro-orbitally in NCD and P-HFD mice and performed a whole-body PET/CT scan (Fig. 6A-B). We then analyzed the biodistribution of radioactivity *ex vivo* of the brain, liver, SWAT, EWAT, and BAT (Fig. 6C-G). PET/CT quantitative analysis revealed that ^18^FDG uptake in the brains of P-HFD fed mice was significantly reduced compared to NCD fed mice (p=0.0068, Fig. 6C). A significant reduction was also observed in EWAT (p=0.0080, Fig. 6D) and in SWAT (p=0.0194, Fig. 6E) following P-HFD, compared to NCD controls. No significant differences were seen in ^18^FDG uptake between NCD and P-HFD mice in the liver (p=0.4011, Fig. 6F), nor in BAT (p=0.1512, Fig. 6G).

**Figure 6.**
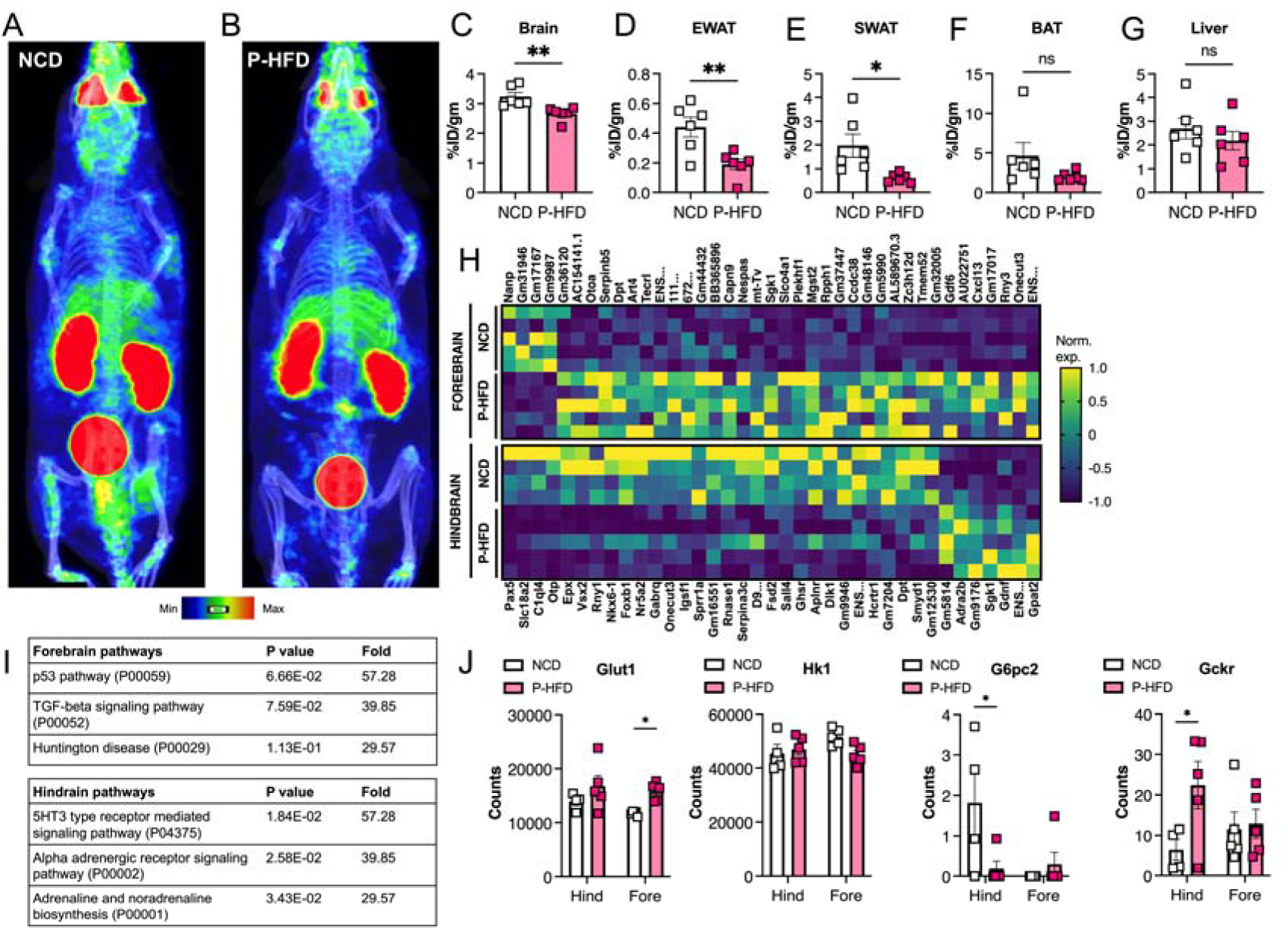
P-HFD induces glucose hypometabolism in the brain and white adipose tissue and influences the brain transcriptome. Representative PET/CT whole-body scan of A. NCD, B. P-HFD mice. Images are displayed according to the intensity scale for tracer activity, from red (highest), green (intermediate), to purple (lowest). Quantitative *ex vivo* analysis of ^18^FDG uptake in C. brain, D. liver, E. SWAT, F. EWAT, and G. BAT of NCD and P-HFD mice. (NCD, n=6, P-HFD, n=6) (*p<0.05, **p<0.01). H. Transcriptomic analyses of the forebrain and hindbrain following NCD of P-HFD feeding (forebrain NCD, n=5, P-HFD, n=5; hindbrain NCD, n=4, P-HFD, n=5). Heatmaps represent differentially expressed genes (p>0,05, +/-1.5-fold change, between diets, per tissue). I. Panther pathway enrichment analysis from forebrain and hindbrain. J. Target gene expression from transcriptomic analysis of genes in glucose metabolism. SWAT, subcutaneous white adipose tissue, EWAT, epididymal white adipose tissue, BAT, brown adipose tissue. Data are mean ± SEM.

We next carried out transcriptomic analysis of the forebrain and hindbrain to assess the impact of P-HFD feeding on gene expression. Differential expression analysis between diets revealed 39 and 37 differentially expressed genes in the forebrain and in the hindbrain, respectively, in response to P-HFD feeding, when compared to NCD (p<0,05, log2FC+/-1.5; Fig. 6H). To functionally classify differentially expressed genes we used the PANTHER pathway enrichment tool (Thomas et al., 2003). There were no significantly enriched pathways in the forebrain, those close to statistical significance (p=0,067, p=0,076), with 39-57 fold-enrichment, represented p53 and TGF-beta signaling pathways, potentially indicating stress or immune activation (Fig. 6H). In the hindbrain, significantly enriched pathways were mainly related to neuronal and neurotransmitter function (e.g., alpha adrenergic receptor signaling) (Fig. 6I). Lastly, we looked at target genes that may be involved in regulating brain glucose metabolism (Fig. 6J), functionally contributing to the phenotypic differences seen following PET/CT imaging (Fig. 6A-B). We found that the glucose transporter Glut1 was increased in the forebrain following P-HFD feeding, compared to NCD-feeding. Expression of the glucose phosphorylating enzyme Hk1 was not altered, however the expression of G6pc2, downstream of Hk1, was decreased in hindbrains upon P-HFD feeding, and near undetectable in forebrains under both conditions. Interestingly, the expression of Gckr, that encodes the glucokinase regulatory protein, also known as Hk4, was increased in hindbrains. Gckr is known to compete with glucose for phosphorylation, thus inhibiting hexokinase action (Van Schaftingen, Detheux, & Veiga da Cunha, 1994). While some components glucose metabolism are dysregulated at the gene expression level, increased expression of Gckr in particular, may explain in part the brain hypometabolism we observed by PET/CT imaging.

## Discussion

Patients with T2D are at greater risk of developing neurocognitive comorbidities such as memory deficits, anxiety, or depression (Deschenes et al., 2023; Hryhorczuk et al., 2013). The metabolic risk factors predisposing to these comorbidities are understudied, in part, due to a lack of models representing discrete stages of metabolic disease. We developed a nutritional model of mild-to-moderate metabolic stress (P-HFD) in which glucose intolerance precedes weight gain. This model impacts specific cognitive parameters in a time-dependent manner. Comparison to reference models of obesity, insulin resistance and T2D (C-HFD, db/db) allows us to attribute specific components of dysmetabolism to decline in specific cognitive parameters (Fig. 7). Glucose intolerance *per se* due to short-term P-HFD feeding induces anxiety-like behaviors. Long-term feeding led to weight gain and recognition memory deficits. *In vivo* imaging revealed decreased glucose metabolism in the brain of P-HFD fed mice, this may contribute to the observed cognitive effects. Here, we provide a novel model for understanding the neurocognitive effects of specific components of metabolic dysregulation (glucose intolerance followed by weight gain) (Fig. 7).

**Figure 7.**
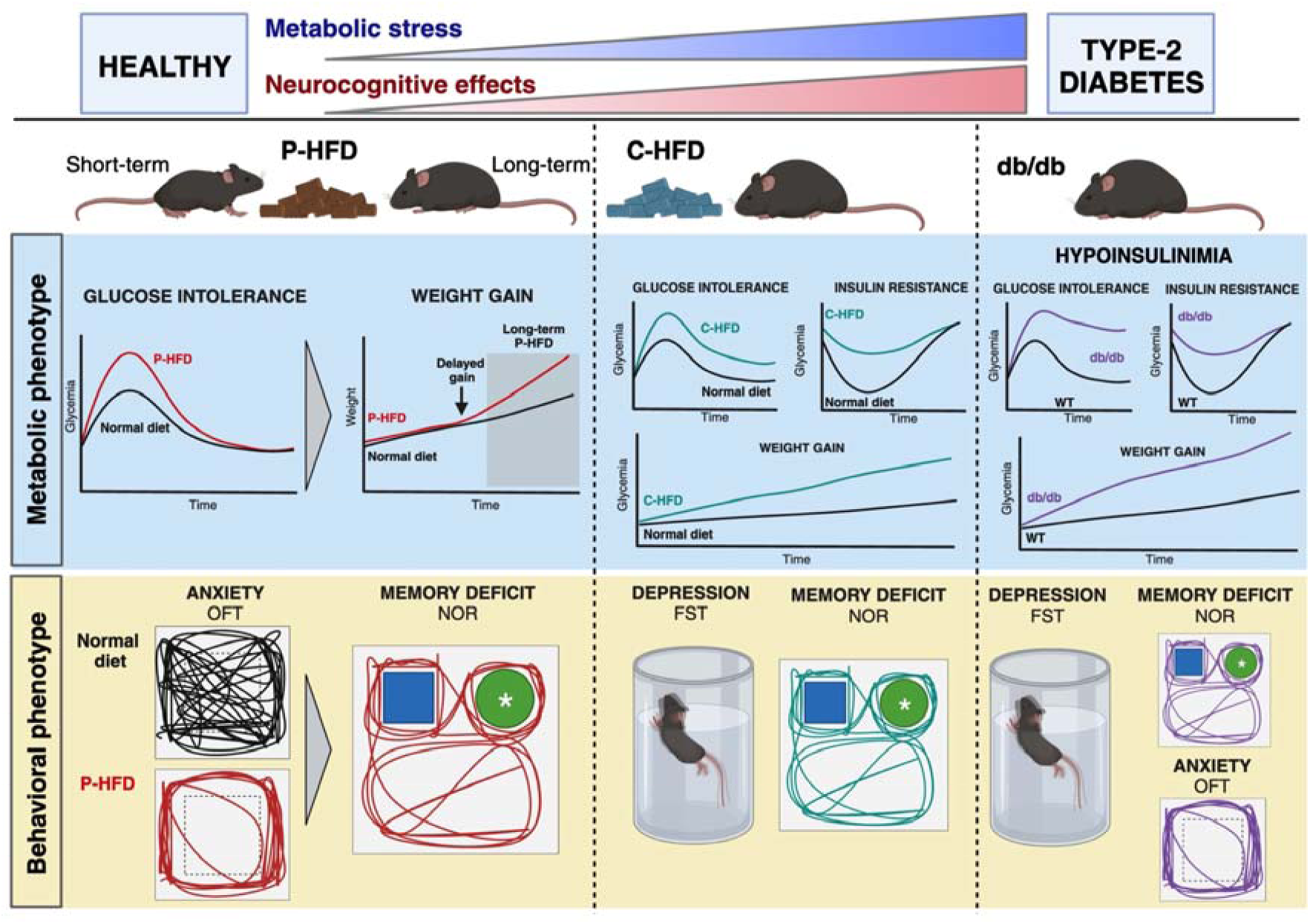
Dysregulation of specific metabolic components is associated with neurocognitive effects. A proprietary high-fat diet (P-HFD) that induces glucose intolerance is associated with anxiety-like behavior in mice, this is followed by weight gain and onset of memory deficits. On a commercial high-fat diet (C-HFD), glucose intolerance, insulin resistance and obesity induce depressive tendencies and memory deficits. A genetic model of T2D (db/db) results in depression, anxiety, and depressive tendencies. (FST: forced swim test; OFT: Open field test; NOR: novel object recognition; WT wild type).

Nutritional analysis of the P-HFD showed that it is moderately high in fat content (42% kcal from fat), placing it in between a NCD control diet (10% kcal from fat) and the commonly used C-HFD (60% kcal from fat). Lower fat content could explain, in part, the delayed weight gain when compared to a C-HFD. However, the lack of a weight difference from the control diet, except after long-term feeding, was unexpected. After analyzing food and water intake in metabolic cages, we did note that food intake was consistently lower in P-HFD fed mice, compared to NCD and C-HFD fed mice. To maintain stable weight, this may have been compensated by increasing fluid intake, since the P-HFD regimen includes 10% (w/v) sucrose in drinking water.

When testing P-HFD for metabolic parameters *in vivo*, we found that insulin sensitivity was unaffected compared to C-HFD mice. A striking outcome of P-HFD feeding was early glucose intolerance, independent of weight gain. This allowed us to attribute specific behavioral outcomes to glucose tolerance alone and to discern its effects from longer-term glucose intolerance and weight gain. For comparison C-HFD-fed mice are obese, glucose intolerant, and insulin resistant within the same timeframe, they develop depressive-like behaviors and recognition memory deficits, but no striking anxiety-like behaviors.

Other studies using the C-HFD nutritional model (60% HFD Research diets D12492, New Brunswick, NJ) reported similar cognitive and behavioral phenotypes, with inconsistent reporting of anxiety-like behavior. Interestingly, the reported phenotypes related to anxiety are revealed following either longer-term administration (16 weeks) (Ogrodnik et al., 2019b) or in mice considerably older than in our cohorts (8 months, versus 5-7 weeks in our study) (Yamada et al., 2011), these additional factors may contribute emergence of anxiety. In the current study, we showed that 13-week C-HFD feeding did not cause overt anxiety, however, total distance travelled was significantly greater than P-HFD-fed and db/db mice in the OFT task, suggesting hyperactivity. Hyperactive behaviors in novel environments and impaired habituation in the OFT have been linked to other neuropsychiatric disorders, such as attention deficit hyperactivity disorder (Zhuang et al., 2001). Other studies show that HFD impacts anxiety-like behaviors in a transient fashion (Gainey et al., 2016), suggesting that both age of animals, when HFD feeding commences, and its duration are key factors in triggering anxiety-like behaviors.

Our previous work showed that db/db mice, which model most features of T2D, including hypoinsulinemia (Burke et al., 2017), have impaired spatial memory associated with thigmotactic behaviors (Al-Onaizi et al., 2022). These are in line with other studies in db/db mice showing anxiety-like behaviors (Dinel et al., 2011; Korolenko et al., 2021; Stranahan et al., 2009). Other evidence shows that db/db mice also display overt depressive-like behaviors (A. N. Sharma, Elased, Garrett, & Lucot, 2010). Here, we confirmed that the db/db mouse model displays anxiety, and depressive-like behaviors, and impaired recognition memory.

Regarding depressive-like behaviors, our data clearly showed that only C-HFD and db/db mice displayed this phenotype, which is in line with previous findings (Abildgaard et al., 2011; S. Sharma & Fulton, 2013; Tsai et al., 2022; Vagena et al., 2019; Yang et al., 2016). P-HFD mice, both at short-and long-term feeding, did not show depressive-like behaviors. These results indicate that insulin resistance, present in C-HFD and db/db models but absent in P-HFD, maybe required for depressive tendencies to manifest. Indeed, insulin resistance has been associated depression and metabolic syndrome (McIntyre et al., 2009; McIntyre et al., 2007). Recent evidence showed the HFD triggers depressive-like behaviors through astrocyte-mediated activation of ventral hippocampal glutamatergic afferents to the nucleus accumbens (Tsai et al., 2022). Use of a selective serotonin uptake inhibitor (SSRI; fluoxetine) alleviated HFD-induced depressive-like behaviors (Tsai et al., 2022). Other studies show that fluoxetine also improves obesity, dyslipidemia, and insulin resistance (Chiu, Tu, Kung, Wu, & Chen, 2021). Hence, insulin resistance appears to be a key modulator of depressive-like behaviors in T2D.

Since we have elucidated which factors of systemic dysmetabolism may be determinants of cognitive and behavioral deficits, we sought to probe potential mechanisms. One important aspect underpinning undesirable neurocognitive outcomes is dysregulated glucose metabolism in the brain. ^18^F-fluorodeoxyglucose (positron emission tomography (^18^FDG-PET) is a method used to evaluate central and peripheral glucose metabolism. ^18^FDG-PET has been widely adopted for clinical evaluation for several neurodegenerative disorders (Minoshima, Cross, Thientunyakit, Foster, & Drzezga, 2022). Indeed, ^18^FDG-PET imaging in a rodent model of Alzheimer’s Disease (AD) showed hypometabolism accompanied by synaptic and neuronal loss (Bouter et al., 2018). Such hypometabolism, observable by ^18^FDG-PET imaging, is increasingly proposed as a unifying marker of neurocognitive dysfunction, including in the context of metabolic diseases such as obesity and T2D (Iozzo & Guzzardi, 2019). We demonstrated that the P-HFD model of glucose intolerance showed robust glucose hypometabolism in key metabolic organs, including the brain. While the dynamics of ^18^FDG PET/CT uptake in neurodegeneration are still not fully understood, findings show that early neurodegeneration is associated with ^18^FDG hypermetabolism, and ^18^FDG hypometabolism in advanced neurodegeneration (Ashraf, Fan, Brooks, & Edison, 2015). Potential mechanisms include transient compensatory reaction in early neurodegeneration that would ultimately lead to exhaustion of vulnerable neural networks, worsening brain pathology. While these mechanisms were not explored in P-HFD mice, our previous work shows that db/db mice display age-dependent neurodegeneration in the hippocampus, a region essential for cognition (Al-Onaizi et al., 2022).

The nutritional model we have developed is of value to scientists in the context of metabolic diseases, namely T2D pathophysiology, and to the field of cognitive and behavioral research. In terms of the diet, more in-depth characterization of lipid composition may yield important mechanistic information, since previous research has found that specific lipid species can modulate mood and behavior (e.g., omega 3 fatty acids)(Liao et al., 2019). Further effects of the P-HFD model on insulin producing and insulin-target tissues can also be characterized. Histological and molecular characterization is indeed required to fully discern the effects on important metabolic organs, such as the liver and adipose tissue, as well as in the brain. Such future work will determine to what extent physiological and adaptive changes in these tissues reflect systemic glucose intolerance. The partitioning of metabolic sequelae using multiple nutritional models can elucidate the determinants of neurocognitive decline, here we demonstrate the glucose intolerance *per se* is associated with anxiety, concurrent weight gain with memory deficits; and that insulin resistance is required for depressive behavior.

## Materials and Methods

### Animals

C57BL/6J (stock number 00664, Jackson Laboratories, Bar Harbor, Maine) were used for this study. At 5-7 weeks old, mice were divided into three groups and were fed either a laboratory-formulated 40% high fat diet (P-HFD), or a commercially sourced 60% high fat diet (C-HFD, Research diets D12492, New Brunswick, NJ), or a normal chow diet (NCD, ssniff, Germany). Only P-HFD received 10% sucrose water, the remaining groups received regular water. Feeding for all groups was *ad libitum*. We used male C57BL/6J-db^-^/db^-^ mice to model type 2 diabetes mellitus (B6.BKS(D)-*Lepr^db^/*J; stock number 00697), and lean C57BL/6J-db^-^/db^+^ mice as controls (stock number 00664; Jackson Laboratories, Bar Harbor, Maine). The animals used for the experiments were the offspring of a breeding colony maintained at the Animal Resources Center of Health Sciences at Kuwait University and the Animal Resource Facility at Dasman Diabetes Institute (DDI). Animals were housed in groups of 3 per cage without environmental enrichment in a temperature-controlled room (12:12 light to dark cycles, 23° C, 50-60% humidity). All mice were weighed, and glucose levels were measured in the tail vein blood using a glucometer (Accu-Check, Roche) at the beginning of the experiment twice every week until the end timepoint of the experiment. At the endpoint of the experiment, animals were sacrificed by transcardial perfusion with saline and the following tissues were harvested and weighed: liver, subcutaneous white adipose tissue epididymal white adipose tissue, and brown adipose tissue. All experiments were conducted in accordance with the regulations of Kuwait University (Animal Resources Center), DDI (Animal Resource Facility) and ARRIVE guidelines. Only male mice were used for all experiments. The protocol for animal studies was reviewed and approved by Animal Ethical Committee of Faculty of Medicine, Health Sciences Center, Kuwait University, Kuwait, 13110 and Animal Ethics Committee of DDI. The experiments were carried out in accordance with recommendations of NIH Guidelines and Guide for the Care and Use of Laboratory animals.

### Diet formulation and diet composition analysis

P-HFD was formulated using hazelnut chocolate spread (35% 155.5g/450g), 100% casein powder (29% 130.7/450g), beef tallow (16% 70.1g/450g), biscuit rusk (17% 75.9g/450g), Stevia rebaudiana derived sweetener (2% 9.1g/450g), vitamin and mineral mix (1% 4.9g/450) and sugar (1% 4.7g/450g). The detailed quantities and composition of the ingredients are reported in Table 1. All dry components were powdered and mixed using a blender. Wet ingredients were mixed with dry ingredients in a stand mixer until all ingredients were combined and homogenous. The mixture was formed and shaped into pellet form and air dried overnight. The laboratory prepared P-HFD was stored in a freezer. Dietary proximate analyses of the P-HFD sample were performed at Kuwait Institute for Scientific Research (KISR) according to AOAC Official Methods. The crude protein-nitrogen was determined using the Kjeldahl procedur*e (American Oil Chemists’ Society (AOAC) official method 2001.11 protein (crude) in animal feed, forage, grain).* The crude fat was measured using the Soxhlet extractor *(AOAC official method 920.39A fat crude in animal feed and pet food)*, and crude dietary fiber and crude ash were determined using the hot extraction method *(AOAC official method 962.09-Fiber crude in animal feed and pet food)*. The Food and Agriculture Organization (FAO) recommendations were used to estimate the carbohydrate content in the P-HFD. The determined fat (F), protein (P), ash (A) and dietary fibers (DF) were used to calculate the amount of carbohydrates [carbohydrates (%) = 100 – (%F + %A + %P + %M + %DF)]. The energy density of the P-HFD was calculated by adding the energy value from proteins, carbohydrates, and fats using the Atwater conversion factor numbers [proteins = 4 kcal/g, carbohydrates = 4 kcal/g, and fats = 9 kcal/g].

**Table 1.** Dietary composition of diets used in the study. Normal chow diet (NCD); Proprietary Western diet (PHFD); Commercial High fat Diet (C-HFD)

|  | NCD | P-HFD | C-HFD |
| --- | --- | --- | --- |
| Crude Fat (g/100g) | 3.9 | 22.55 | 35 |
| Crude Protein (g/100g) | 14.3 | 21.14 | 26 |
| Crude Carbohydrates (g/100g) | 59.3 | 50.21 | 26 |
| Crude Fiber (g/100g) | 4.2 | 3.93 | 6.46 |
| Crude Ash (g/100g) | 5.0 | 2.17 | N.A |
| Total kcal/g | 3.35 | 4.88 | 5.24 |
| Calories from fat (%) | 10.478 | 41.558 | 60 |
| Calories from protein (%) | 17.075 | 17.315 | 20 |
| Calories from carbohydrates (%) | 72.447 | 41.126 | 20 |

### Glucose and insulin tolerance tests

For the oral glucose tolerance test (GTT) and insulin tolerance test (ITT), mice were fasted for 6 hours, and their weights were measured. For both tests, the fasting glucose levels were measured at t=0 via tail vein blood using a glucometer (Accu-Check, Roche). For OGTT, 2g/kg dosage of glucose solution (20% D-glucose in sterile 0.9% NaCl saline solution) was orally administered using flexible plastic feeding tubes with time lapse of 2-3 minutes between each animal. For ITT, an intraperitoneal injection of insulin (Actrapid, UK) at a dose of 0.5UI/kg, or 0.75UI/kg (for P-HFD and C-HFD) was administered. For both OGTT and ITT, glucose measurements were measured at 15, 30-, 60-, 90-, and 120-mins post administration.

### Metabolic cages

Mice were housed individually in metabolic cages at weeks 1, 3, and 8 of the experiment. The Promethion High-Definition Behavioral Phenotyping System (Sable Systems International) was used to metabolically characterize mice. This system consisted of a two-door configuration for 16 mouse cages. This Promethion Environmental Sensor Assay (ESA) integrated temperature, light, humidity, barometric pressure, and enclosure activity with metabolic and behavioral data. The chambers were linked to an open-circuit flow-through calorimetric device connected to a computer-controlled system of data acquisition. Mice were habituated in the metabolic cages 24 h before measurement. Oxygen consumption (VO2) and carbon dioxide production were measured every 3 min for 24 h. Measurements were recorded and divided during between day 1 (0800 and 1959), night (2000 and 0359), and day 2 (0400 and 0759). During 24 h the respiratory quotient (carbon dioxide production-to-VO2 ratio), the average energy expenditure, food intake and water consumption were measured and recorded. Data was available real-time via the browser-based data management application.

### Novel object recognition task

Mice were acclimatized to the experimental room for 1 hr prior to testing. The test was performed in an open field box (WxLxD= 50cmx50cmx50cm). During the learning phase, mice were placed in presence of two identical objects and were allowed to freely explore the objects for 5 minutes, after which they returned to their home cages. The box and objects were cleaned with 70% ethanol immediately before and at the end of each testing animal to minimize olfactory cues. Mice were re-introduced to the chamber 1 hour later for the recall phase. Mice were left for 5 minutes in the presence of one of the familiar objects along with a new object with similar size, texture, and shape. A digital camera was placed above the open field and connected to a computer with a video-tracking system (ANY-Maze, Stoelting Co, IL, USA) which objectively measures and records parameters such as time spent exploring the objects. Object exploration was defined as either touching or exploring the object with less than 5 cm proximity. The experimental box and the different objects were systemically cleaned between each test and each mouse. Mice were randomized and mice whose exploration time is less than 20 seconds during the training sessions are excluded from analysis. The total exploration time was calculated Recognition Index (RI) = time novel (TN)/(TN + time familiar (TF)) and Discrimination Index (DI) = (TN − TF)/(TN + TF) were analyzed.

### Open field test

Open field test is used to analyze locomotion and anxiety-like behavior using a sound insulated open field box (WxLxD= 50cmx50cmx50cm) with a digital camera oriented above the open field and connected to a computer with a video-tracking system (ANY-Maze, Stoelting Co, IL, USA) which objectively measures and records movements. Mice were acclimatized to the experimental room for 1 hr before being introduced into the chamber. Mice were allowed to freely move in the novel environment for 30 minutes. Their movements in the box during were recorded. The distance travelled, number of entries and time spent in the peripheral and central zone were assessed.

### Forced Swim Test

Mice were subjected to the forced swim test to assess depressive-like behaviors. Mice were placed in the testing room to acclimate to the environment for 1 hr before testing. Next, each mouse was placed in a 2 L beaker with 1.8 L of water (25-27°C) for 6 minutes. A video camera was placed above the beaker and immobility time, immobility episodes and distance swam were recorded and analyzed using ANYMaze software (Stoelting Co., IL). The first minute of activity was not used for the analysis, results were presented as data from 2-6 minutes testing. Water was changed after every 2 mice completed testing.

### Morris Water Maze

This task was performed as previously described (Al-Onaizi et al., 2022). Briefly, mice were given two 90s sessions per day for training for four consecutive days, with a 5-minute inter-trial interval. Mice that did not reach the platform during the training sessions were gently guided to the platform. On the fifth day, memory was tested with a probe trial (one 60s session). In the probe trial, the platform was removed, and time spent in each quadrant of the pool was measured and calculated. The experiment was performed in a 95-cm diameter pool filled with 18-20°C water. The platform was submerged one cm below the water surface, and different spatial cues were distributed around the pool. The mice were acclimated to the test room for one hour prior to the experiment for each day of the experiment. Sessions were recorded and data were analyzed using a video tracking software (ANY-Maze, Stoelting Co, IL, USA). Parameters of spatial learning performance, such as latency to reach the hidden platform and time spent in quadrant were recorded and analyzed.

### In Vivo PET/CT Imaging and biodistribution analysis

All animal experiments were conducted in accordance with Kuwait University Animal Resources Center approved protocol. Mice fasted for at least 6 hours. They were anesthetized with ketamine (100mg/kg) and xylazine (25mg/kg) in 0.9% sodium chloride and received intra-orbital injection of clinical grade ^18^F-FDG (127 ± 20 µCi; 4.7 ± 0.7 MBq). Mice were kept under anesthesia for 60 min post injection then a whole-body PET scan (NanoScan® SPECT/CT/PET) was obtained using a 1:5 coincidence mode (where each detector module is in coincidence with five opposing ones and is equivalent to maximum field of view of 212_×_212_×_283 mm). Then, a CT scan was acquired (semi-circular full trajectory, maximum field of view, 360 projections, 35 kVp, 170 ms and 1:4 binning) for attenuation correction. For image reconstruction (whole-body Tera-Tomo 3-dimensional reconstruction algorithm and the following settings: 4 iterations, 6 subsets, full detector model, low regularization, spike filter on, voxel size 0.4 mm and 400–600 keV energy window. PET data were corrected for randoms, scatter and attenuation. InterViewFusion software (version 3.09.008.0000) provided by Mediso was used to analyze the reconstructed images. Mice were kept under anesthesia for 60 min post injection then were euthanized for biodistribution analysis. Blood was collected by cardiac puncture. Major organs (liver, brain, SWAT, EWAT and BAT) were harvested, weighed, and counted in an automated γ counter (1470 Wizard, PerkinElmer, USA). The percent injected dose per gram (% ID/g) of tissue and the percent injected dose per organ (%ID/organ) were calculated by comparison with standards of known radioactivity.

### RNA-seq

RNA was extracted from tissues using Qiagen RNeasy mini kit (Qiagen, Hilden, Germany) and prepared for Illumina sequencing as previously described (Dashti et al., 2022). Raw RNA-seq data were preprocessed using Trimmomatic (version 0.39) to remove low-quality bases and adapter sequences. Paired-end reads were quality trimmed and filtered. The quality-trimmed reads were aligned to the reference genome using HISAT2 (version 2.1.0). Aligned SAM files were processed using HTSeq (version 0.6.1p1) to quantify reads mapping to gene features.

The count matrix obtained from HTSeq was used to create a DESeq2 dataset. Sample metadata, including experimental conditions and replicates, were incorporated into the analysis. DESeq2 models were fit to the dataset to detect differentially expressed genes between NCD and P-HFD

### Statistical Analysis

All data were presented as mean ± SEM. Shapiro-Wilks test was used to check for the normality of data. When comparing two sample means, Student’s t-test was used. A one-way or two-way analysis of variance (ANOVA) with repeated measures (RM) was performed when comparing more than two sample means. When appropriate, a Bonferroni post hoc analysis test was used. GraphPad Prism (v10.0) was used for all statistical analysis. The test results were considered statistically significant at the P < 0.05 level.

## Author statement

All authors have read and approved the final version of the manuscript. The authors declare no conflict of interest and contributed to the study as follow; MA-O: conceptualization, investigation, formal analysis, writing original draft, writing review editing, funding acquisition; KB: investigation, formal analysis; SK: formal analysis, writing review editing; DAlt: formal analysis, writing review editing; SD: investigation; DAla: investigation; HA: investigation; MA: investigation; AK: investigation; RN: investigation; MRW: investigation; RA: formal analysis, writing review editing; HA: formal analysis, writing review editing; FA-M: writing review editing, funding acquisition; FAlz: investigation, formal analysis, writing original draft, funding acquisition.

## Acknowledgements

This work is supported and funded by Kuwait University Research Grant No. RM01/19 (to M.A-O) and Research Core Facility Grant No. SRUL02/13. FA is supported by KFAS grants RA AM-2022-009 and RA AM-2023-007. We thank the Animal Resources Center (Kuwait University) staff for technical support. We would like to thank the Nuclear Medicine Department staff at Mubarak Al-Kabeer Hospital, Ministry of Health, State of Kuwait, for providing ^18^F-FDG. The authors would like to extend their gratitude for PET/CT (Research Grant No., GM18/01) and biodistribution technical assistance to Mrs. Heba Essam Abd Elghany, Ms. Fatma Dashti, Dr. Manal Abdallah Morsy and Dr. Mohammad Sakr.

